# Imaging of sub-cellular fluctuations provides a rapid way to observe bacterial viability and response to antibiotics

**DOI:** 10.1101/460139

**Authors:** Charlotte R Bermingham, Isabel Murillo, Alexandre D J Payot, Krishna C Balram, Maximilian B Kloucek, Simon Hanna, Niamh M Redmond, Helen Baxter, Ruth Oulton, Matthew B Avison, Massimo Antognozzi

## Abstract

Determining the viability of bacteria in a sample is an essential microbiological technique used in healthcare, industrial bioprocesses and research. Increasingly, attention has been focussing on antimicrobial susceptibility tests (AST), allowing rapid and appropriate prescribing of antibiotics. Current AST are limited in speed as they rely on detecting growth of microorganisms. Faster AST could be enabled by the recent discovery that living bacteria manifest nano-scale fluctuations, which reduce when the bacteria die. Here, we demonstrate a direct method of visualising fluctuations within bacterial cells using Sub-Cellular Fluctuation Imaging (SCFI), which is based on Total Internal Reflection Microscopy (TIRM). We show that SCFI can measure the viability of bacterial samples within minutes, distinguishing not only between live and dead bacteria but also live bacteria in different metabolic states. Importantly, we subsequently show that SCFI can rapidly distinguish antibiotic-treated resistant and susceptible bacteria, and therefore has particular application as a rapid AST.

## Introduction

It is well established that current ASTs typical take 18 hours using traditional petri dish or broth-based growth detection methods with up to 36 hours to prepare pure bacterial cultures for the test (Jorgensen and Ferraro, 2009). The waiting time in obtaining AST results can mean that, for several days, patients receive ineffective antibiotics, potentially resulting in prolonged symptoms, complications, fatalities and increased costs to the healthcare system (Barenfanger et al., 1999). In addition, unnecessary and broad-spectrum antibiotics are often prescribed empirically, which contributes to the alarming rise of antimicrobial resistance (Goossens et al., 2005). Automated AST and techniques detecting early stage growth can reduce the duration of these tests to several hours from cultured bacteria (Marschal et al., 2017). The fundamental limitation is the time required for bacteria in the sample to replicate to a sufficient density to allow detection. Genotypic tests such as Polymerase Chain Reaction (PCR) based methods designed to identify resistant bacteria, even directly from clinical samples, can be rapid, however, prior knowledge of the resistance mechanism is needed, limiting their use to specific cases such as Meticillin Resistant Staphylococcus aureus (MRSA) detection (Dinarelli et al., 2015). The development of rapid AST is vital to enable informed prescribing at the earliest stage (Okeke et al., 2011) and is recognized by the World Health Organisation as being of great importance to combat antimicrobial resistance (WHO, 2015).

It has been previously discovered using Atomic Force Microscopy that live cells exhibit nanoscale fluctuations related to their metabolism (Pelling et al., 2004). More recently, it has been observed that viable bacteria also exhibit nanoscale fluctuations, while dead ones do not (Longo et al., 2013), (Etayash et al., 2016), (Syal et al., 2015), (Johnson et al., 2017). This discovery provides an alternative bacterial phenotype that can be measured to perform viability and AST tests that are no longer limited by the requirement for bacteria to replicate up to a certain threshold density for detection. So far, the nano-fluctuations of bacteria have been detected using micro-cantilevers decorated with bacteria (Longo et al., 2013), a quartz microbalance (Johnson et al., 2017) and plasmon resonance (Syal et al., 2015). These techniques demonstrate the potential of using fluctuation assays to discern the viability of bacteria in a sample. However, these methods cannot locate the actual source of the fluctuations, making it difficult to interpret the results and to speculate on the underlying biochemical process. Furthermore, the complexity of some of these methods could obstruct their widespread use in healthcare settings.

Here we present a label-free optical method, termed Sub-Cellular Fluctuations Imaging (SCFI), which enables the direct observation of fluctuations with sub-bacterial spatial resolution. SCFI can quantify the magnitude and location of these fluctuations to determine the state of individual bacteria in a few seconds. Bacteria are bound to a glass surface using antibodies, and illuminated using an evanescent field (Figure 1A). From the diagram it is clear that only a small portion of the bacterium is illuminated and therefore SCFI is sensitive to changes occurring only within this narrow region. A more detailed description can be obtained by modelling the optical interaction between an evanescent wave and a bacterium deposited on the glass-water interface (Figure 1B). The model uses the finite-difference time-domain method (FDTD) which involves solving the wave equation on a fine grid of points superimposed to Figure 1A. The model shows that the evanescent field decays by more than half of its value at the glass surface within 100 nm. If we now focus our attention to the region of the bacterium within the evanescent field (Figure 1C) we find that the space between the outer and inner bacterial membranes contains several proteins that could cause local fluctuations, resulting in changes of the local scattering of the evanescent field. Scattering of the evanescent field in this region will be significantly larger than scattering further away from the inner membrane.

**Figure 1.**
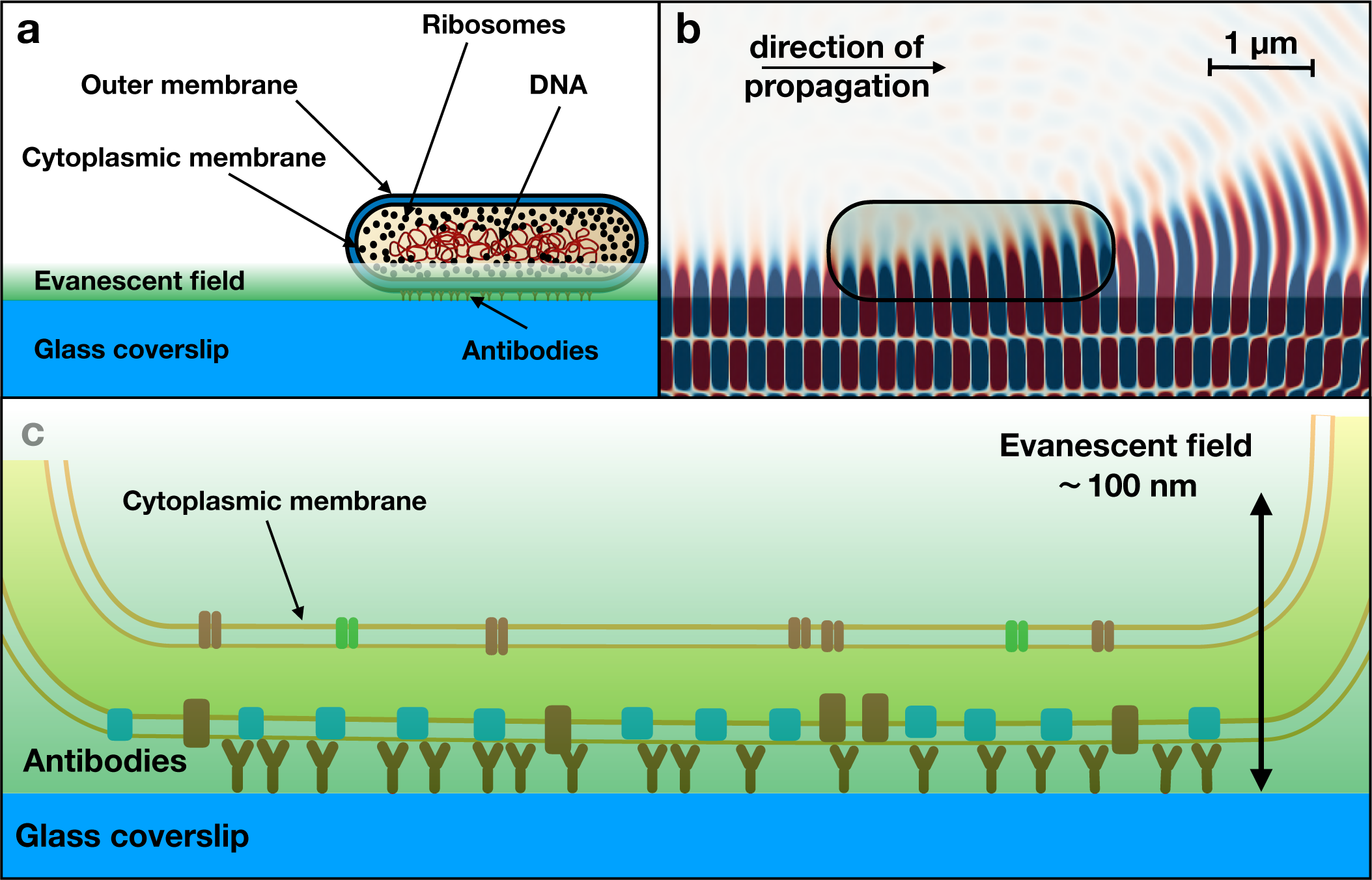
The interaction between an *E. coli* bacterium and the evanescent field. (**A**) Diagram of an *E. coli* bacterium with its main components and illuminated using SCFI. Only the lower part of the bacterium is in the evanescent field generated by total internal reflection of a laser beam. (**B**) Finite-difference time-domain (FDTD) simulation of the electromagnetic field generated using SCFI at the glass-water interface with a bacterium. The model shows how part of the evanescent wave is frustrated by the bacterium and propagates in the water. The model confirms a small penetration in the cell. At 100 nm from the glass surface the value of the field is less than half its value at the surface. (**C**) Close-up model of the region of the bacterium illuminated by the most intense part of the field, including the tethering antibodies. Most of the scattering collected by the video camera is originated in this area.

The optical set up used in SCFI (Figure 2A) is based on the Scattered Evanescent Wave (SEW) detection system, originally designed to detect micro-cantilever tips for vertical probe microscopy (Antognozzi et al., 2008), (Antognozzi et al., 2016). A high numerical aperture objective lens is used to generate the total internal reflection of a laser beam and to collect the light scattered by the sample. A video camera produces a high-resolution image of the bacteria when illuminated by the evanescent field. The microscope can also operate in the more conventional bright field illumination to obtain standard optical images of the bacteria, such as *E. coli* (Figure 2B) in addition to the images produced from the scattered evanescent light (Figure 2C). The bright spots at the poles of the bacterium are typical of the scattering pattern produced by elongated objects aligned in the direction of propagation of the evanescent field (Figure 2 - figure supplement 1). Bacteria in different alignments relative to the evanescent field produce different scattering patterns (Figure 2 figure supplement 2). Bacteria were aligned in the direction of propagation of the evanescent field using a flow cell so that the data is consistent. Real-time recordings using SCFI of the scattered light from a bacterium show intensity changes within the bacterial envelope that have their origin in nano-scale movements as no movement is observable in bright 1eld microscopy.

**Figure 2.**
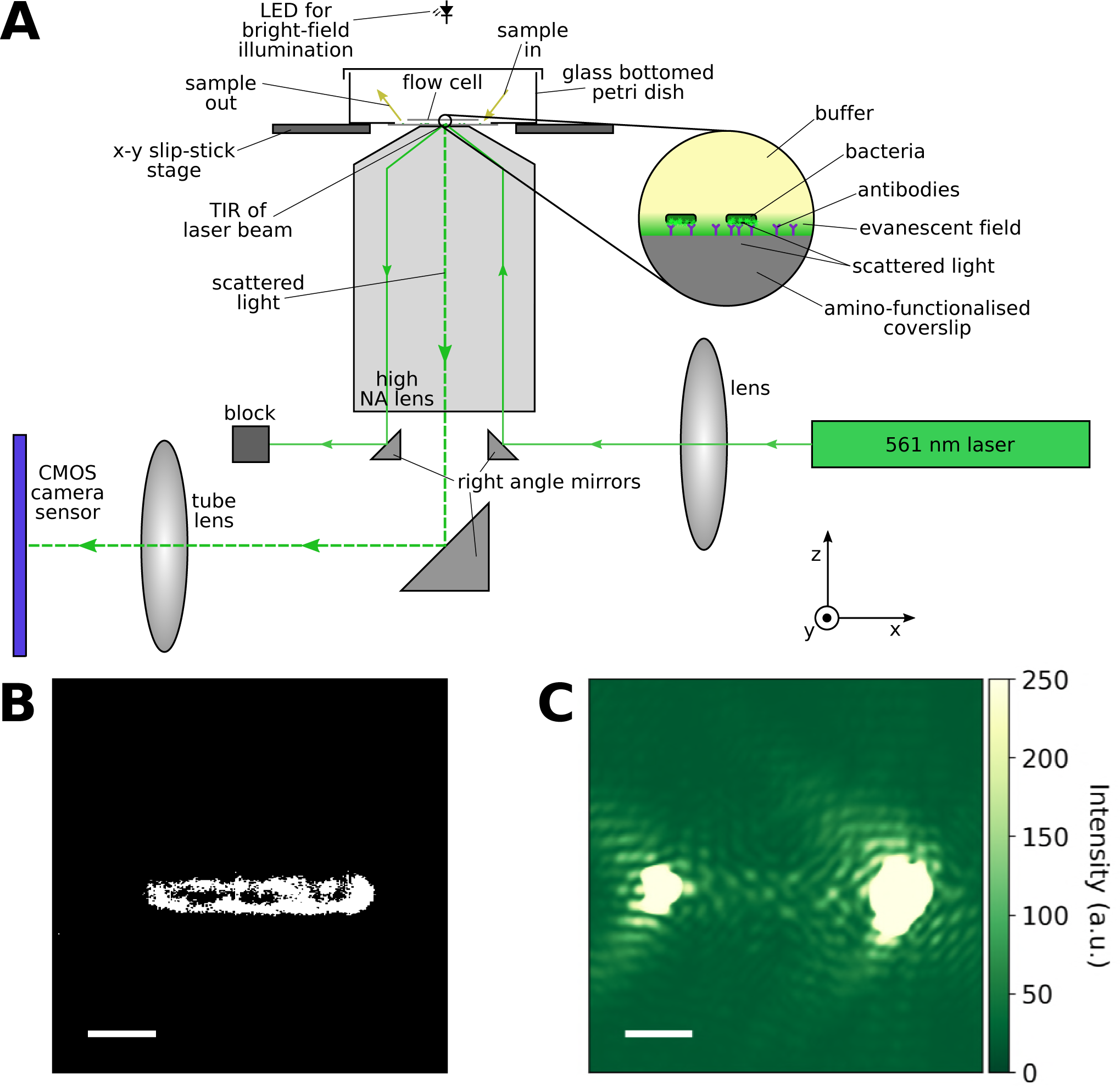
Details of the scattered evanescent wave detection system. (**A**) Schematic diagram of the set up. A laser beam is totally internally reflected at the sample surface to generate an evanescent field decaying away from the surface. Bacteria are bound to the surface in a flow cell via antibody attachment. Scattered light from the surfaces of bacteria within the evanescent field is collected and focused onto a high resolution CMOS camera in order to image nanoscale fluctuations within the bacteria. The flow cell is contained within a functionalised, glass bottomed petri dish and sits on an x-y slip-stick stage for horizontal positioning while a flexure mechanism provides vertical movement for focussing. (B) Standard optical microscopy image of a single E. coli bound to the coverslip surface, imaged in our system. (C), Scattered evanescent wave image of the same bacterium imaged in our system. The scale bars are 1*μ*m. The optical image shows the shape of the bacterium but no features at the sub-bacterium level, whereas the scattered light image displays variation within the bacterium. These intensity variations fluctuate in time as the cell fluctuates. **Figure 2 - figure supplement 1**: Modelling of the scattering pattern. The scattering pattern produced using finite element modelling for bacteria aligned with the field. The two intensity peaks observed experimentally at the ends of the bacteria can be seen. **Figure supplement 2**. Scatter patterns produced by bacteria in different orientations relative to the evanescent field propagation direction. (A) Bacteria aligned with the direction of the propagation of the evanescent field, showing strong scattering at the poles of the bacterium which saturates the camera. (B) Bacteria perpendicular to the direction of propagation of the field, showing strong scattering down the length of the bacterium on either side. The red arrow indicates the evanescent field propagation direction. In order to take consistent data, we aligned the bacteria using a flow cell and only took measurements from bacteria that are well aligned with the field.

## Results

### Determining the metabolic state of bacterial samples using SCFI

In order to determine if SCFI can discriminate between bacteria in different metabolic states, three different sample preparations were designed to produce the same *E. coli* strain but with different levels of metabolic activity. *E. coli* was selected due to the clinical relevance of this species, being, amongst other things, the most frequent cause of urinary tract infections (UTIs) (Flores-mireles et al., 2015), which are the most common and burdensome community and hospital acquired infection (Wilson and Gaido, 2018), (Ahmed et al., 2018). All samples were prepared from an overnight culture in Lysogeny broth (LB). One sample, which we will refer to as dead, was subcultured in fresh LB to an Optical Density at 600 nm (OD600) of ~ 1 then killed using 2% formaldehyde in PBS and resuspended in 0.06% sodium azide in PBS to prevent contamination with living cells.

A second sample (named exponential) was subcultured in fresh LB to an OD600 of 0.205, which we confirmed experimentally to be in the exponentially growing state (Figure 4 - figure supplement 1), in agreement with literature values (Sezonov and Ari, 2007). A third sample (named stationary) was taken directly from the overnight culture and had an OD600 of 4.44, which we confirmed to be in a stationary state (Figure 4 - figure supplement 1). Bacteria from the three different samples were aligned and bound on different glass coverslips using antibodies according to the protocol described in the Methods.

To gain an initial appreciation of the difference between the SCFI recordings of bacteria in different states the exponential sample (Figure 3A and B) is compared to the dead sample (Figure 3C and D). The fluctuations of one bacterium were recorded as a sequence of frames and displayed as a heat map (Figure 3B and D). Each pixel of the map represents the standard deviation of the intensity for that pixel during a 20 second recording. The bacterium from the live, exponential sample (Figure 3B) shows a much higher level of fluctuation than the bacterium from the dead sample in (Figure 3D) confirming that SCFI can readily differentiate between these two bacterial states. The original videos are shown in video 1 and video 2 for the live and dead data respectively.

**Figure 3.**
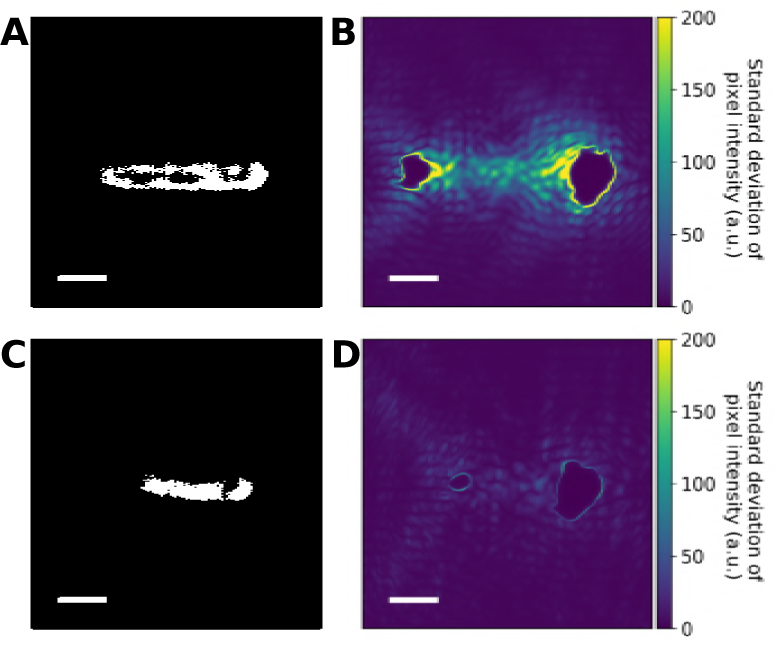
The fluctuation levels of live and dead bacteria. (**A, C**) Optical images of a live (**A**) and dead (**C**) bacterium. (**B, D**) Heat maps displaying the standard deviation in intensity of each pixel in a 20 s video of the live (**B**) and dead (**D**) bacteria in A and C taken using SCFI. The live bacterium exhibits significantly higher intensity fluctuations corresponding to much greater sub-bacterium movements than the dead bacterium, which is almost motionless. The poles of the bacteria show no variation due to saturation of the camera detector. The original videos are video 1 and 2 for the live and dead bacteria respectively.

To determine if SCFI can provide statistically significant differences when analysing a large number of bacteria in these three states, we recorded the fluctuation signals of approximately 50 bacteria from each sample. In order to quantify the level of fluctuation for each bacterium, a 20 s video of each bacterium was recorded at 20 frames/sec using SCFI. An area of 30 by 30 pixels centred on the bacterium is used for analysis and the pixel intensities are normalised by the average intensity of the analysis region in the video. For each pixel in the analysis area, the standard deviation of its intensity through the frames in the video is calculated. The average pixel standard deviation in the analysis area, σ, is then calculated to quantify the fluctuation level of the bacterium (Figure 4 - figure supplement 2). Representative distributions of the *σ* values for the three samples are displayed in Figure 4.

**Figure 4.**
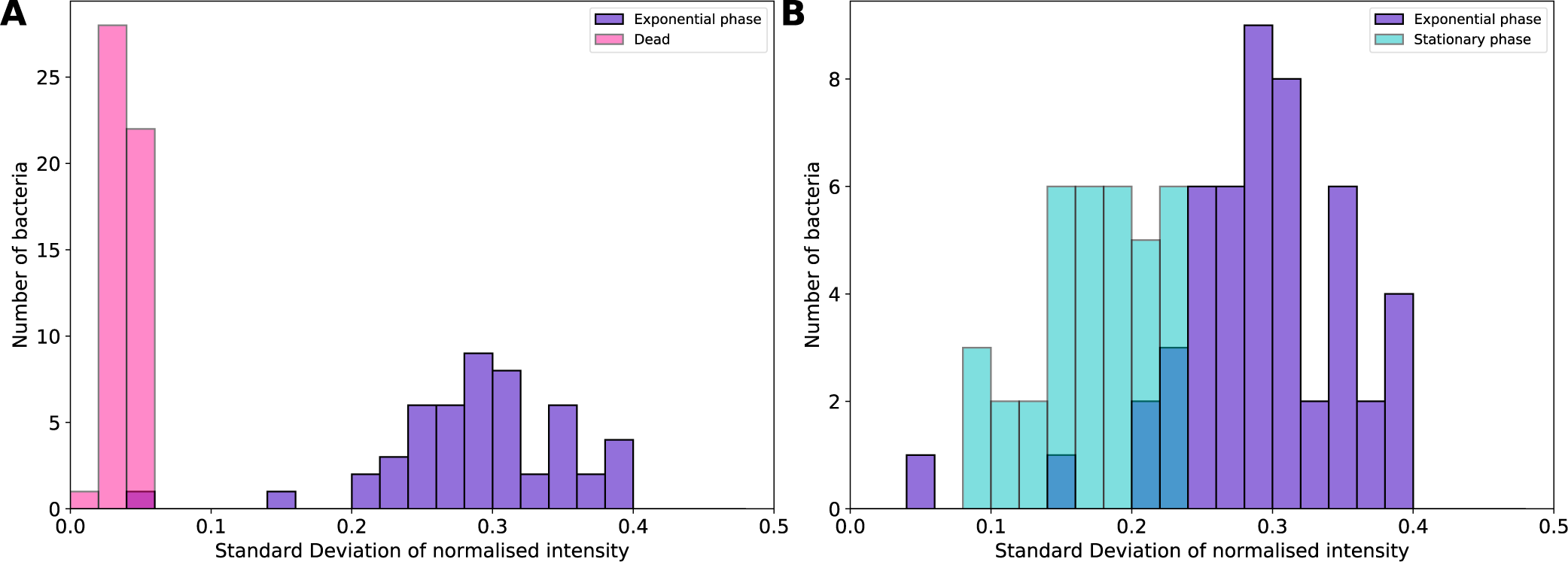
Measurements of the fluctuations of bacteria in different metabolic states. (**A**) The distribution of the fluctuation levels for exponentially growing bacteria (n=50) and dead bacteria that have been killed using formaldehyde (n=51). The exponential bacterial sample displays a wide range of fluctuation levels whereas the dead bacteria have consistently low fluctuation levels. Comparing the distributions with the t-test yields a p-value < 0.001. (**B**) The distribution of the fluctuation levels for exponentially growing bacteria (as in **A**) compared to live bacteria in a stationary state (n=36). There is a clear difference in the mean fluctuation level, with a p-value < 0.001. This data shows the effectiveness of our technique in distinguishing between bacterial samples in different metabolic or fitness states. **Figure supplement 1.** Growth of E. coli. The OD600 of an overnight suspension of DH1 E. coli grown at 37 degrees in a shaker (overnight suspension) and of a subculture in fresh LB taken from the overnight suspension (inoculation). The lag phase of approx. 20 minutes, exponential growth phase up to about OD600 0.4 and slowing growth of the subculture can be seen. The overnight culture is in a stationary state. The OD600 values of the stationary and exponential state samples used in Figure 4 are indicated. **Figure supplement 2.** The data analysis process. (**A**) A bright-f eld image of the bacterium is used to locate the centre of the bacterium. (**B**) This is then used to locate a 30×30 pixel area in the centre of the bacterium to be analysed in each frame in the scattered light video. (**C**) Normalised pixel intensities for a live bacteria. The intensity of each pixel in the 30×30 region of interest is extracted from the video, the values are normalised by the mean intensity of the 30×30xnF analysis region (nF the number of frames). Some of these pixel intensities are plotted (full coloured and grey lines) to illustrate how they fluctuate throughout the video for a live bacteria. Long period oscillations (period>6 sec, frequency<0.17Hz) are removed from the pixel intensities by subtracting the rolling average using a triangular window of width 9 secs (180 frames) (full black lines). The standard deviation of each pixel’s normalised and f ltered intensity is then taken (coloured dashed lines) and averaged over the 30×30 region of interest producing the final fluctuation measure. (**D**) Normalised pixel intensities for a dead bacteria. Three intensity traces for dead bacteria are highlighted for comparison with the live traces. They show much lower fluctuation levels for the dead bacteria. **Figure supplement 3.** Comparing the fluctuation distributions of identical samples throughout the day. The fluctuation distributions for two samples prepared in LB from separate subcultures from the same overnight culture such that they are exponentially growing were compared to assess reproducibility. The early sample (n=51) was made up and imaged approx. 9 hours before the late sample (n=52). The distributions are statistically similar (p-value = 0.4) and have identical means (0.12 +/− 0.03), therefore samples from different subcultures made on the same day from the same overnight culture can be reliably compared.

Figure 4a shows that there is a clear difference in the distribution of fluctuation levels for the dead and the exponential samples. The average *σ* of the exponential bacteria (0.29 ± 0.06 (s.d.)) is approximately 5 times that of the dead bacteria (0.038 ± 0.008). Interestingly, bacteria in the dead sample have a much smaller spread of fluctuation values than the exponential sample. Considering that the exponential sample contains bacteria in different stages of the cell life cycle it is likely that this heterogeneity is reflected in the range of fluctuations recorded. We hypothesise, therefore, thatthe spread of fluctuations seen in a sample is an intrinsic characteristic of a particular sample and not just experimental noise.

To further confirm this hypothesis, we compared the distribution of fluctuations in the exponential and stationary samples (Figure 4B) as they contain bacteria in different metabolic states. Using SCFI we found that the fluctuation distribution for the stationary sample has a mean of 0.17 ± 0.04 that is statistically significantly different from the exponential sample (p-value ≪ 0.001, t-test). Differently from Figure 4A the distributions in Figure 4B have a large overlapping region strongly suggesting that the exponential sample contains bacteria in a similar metabolic state to those in the stationary sample. The measurement of metabolic states according to their *σ* distribution is unique to SCFI and is a clear indication of how SCFI can provide a much more informative viability test. In particular, it can differentiate samples according to their different metabolic activities and it can reveal variations within a population, giving a more complete picture of the sample state. As a control to check the reproducibility of measurements, two exponential samples were grown and analysed at the beginning and end of a day of measurements and produced statistically similar results (Figure 4 - figure supplement 3).

These preliminary results clearly indicate that SCFI could provide not only a viability test, but a rapid fitness test or cell “activity ”test based on an absolute scale of bacterial activity. As well as having considerable value in infectious diseases diagnostics, this has important implications for the biotechnology industry where continuous monitoring of the health of bacterial cultures is crucial. Potentially, SCFI can be used to test the viability of unculturable bacteria that cannot be investigated using growth based methods or methods requiring dense, pure cultures.

### Determining the susceptibility of bacterial samples to antibiotics using SCFI

To assess the effectiveness of SCFI when testing antimicrobial susceptibility, we measured the effects of the bactericidal antibiotic kanamycin on an ***E. coli*** strain that was susceptible to kanamycin and on one that was transformed to kanamycin resistance by introduction of a plasmid. Kanamycin is an aminoglycoside, therefore it works by binding to the 30S ribosomal subunit resulting in misreading of mRNAso the bacterium cannot properly synthesize proteins for growth and rapidly dies as a result of the build-up of misfolded proteins. The two ***E. coli*** samples were grown to exponential state then incubated in a 37 °C shaker for 30 mins in LB containing 100 *μ*g/ml kanamycin along with control samples in LB containing no kanamycin. They were then resuspended in LB without antibiotic for imaging. Plating and incubating the resulting suspensions on agar showed that 99.9997% of the susceptible bacteria were rendered non viable whereas the resistant ***E. coli*** population was as viable as if not treated with antibiotic.

The SCFI fluctuation distributions were measured for ***E. coli*** samples that are susceptible (Figure 5A) and resistant (Figure 5B) to kanamycin. Repeat measurements were also taken (Figure 5 - figure supplement 1). For the kanamycin susceptible strain there is a statistically significant difference (p-value ≪ 0.001, t-test) in the distributions of σ between the untreated and kanamycin-treated ***E. coli*** (Figure 5A). In contrast, there is a much lower statistically significant difference (p-value = 0. 015, t-test) between the untreated and kanamycin-treated kanamycin resistant ***E. coli*** (Figure 5B). Previous findings by Longo et. al. (Longo et al., 2013), observed that the fluctuations of resistant bacteria reduced on introduction of an antibiotic, but recovered gradually over 15-20 minutes to the pre-antibiotic fluctuation levels, implying that the bacteria were affected but not killed by the antibiotic. SCFI measurements confirm that kanamycin resistant bacteria have only slightly lower fluctuations than untreated samples after 30 minutes exposure to the antibiotic. The slight difference could be due to the antibiotic compromising the bacterial health while not killing them. Differently from reference 9, we do not inject antibiotics directly into the sample to detect relative differences but treated and untreated samples are observed in the same buffer conditions allowing a fairer comparison between the two populations. Furthermore, this protocol is more similar to a standard diagnostic test in which an untreated sample (control) is compared to a treated sample in similar conditions.

**Figure 5.**
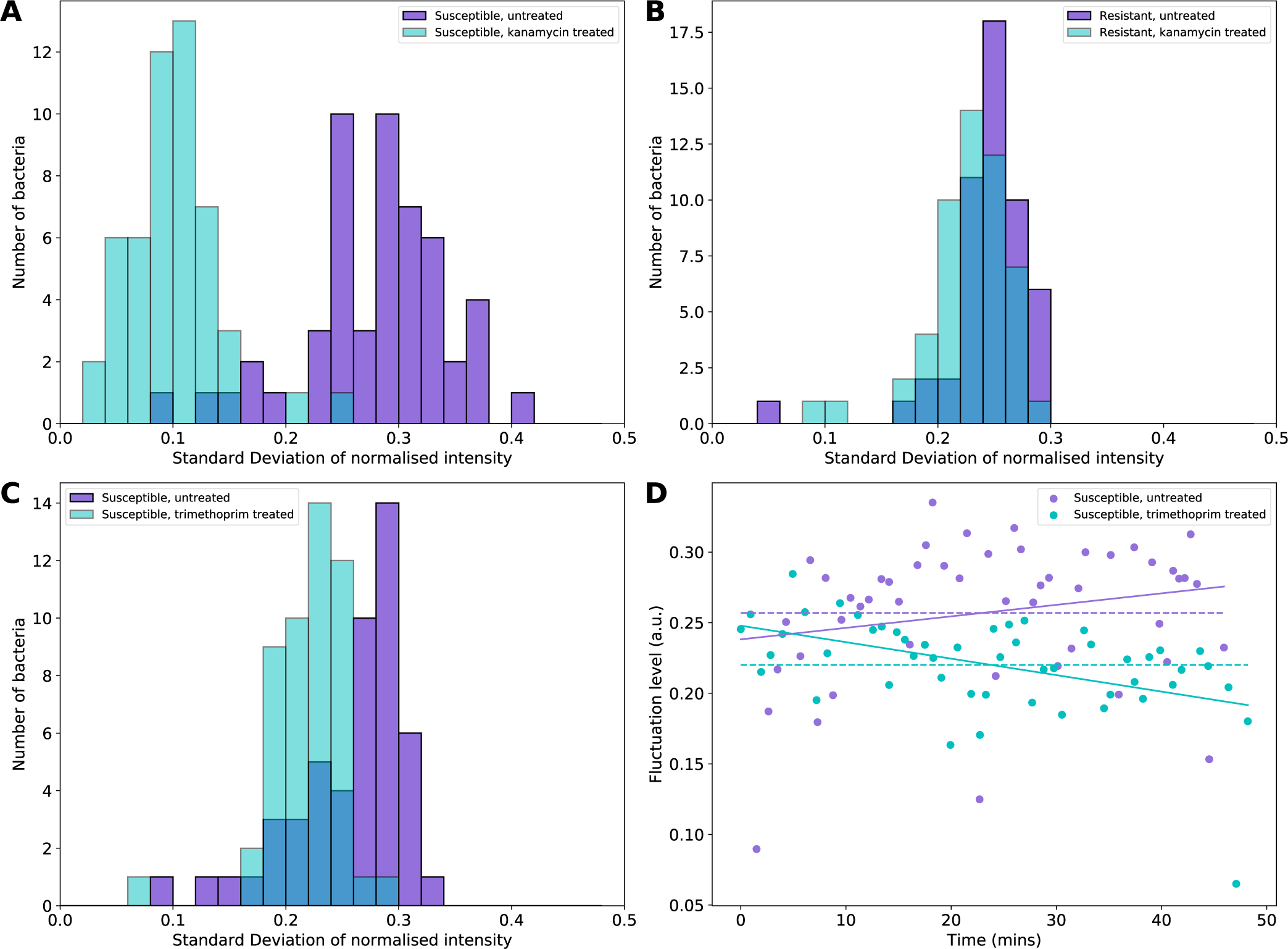
(**A**) The distribution of the fluctuation levels for untreated and kanamycin treated susceptible bacteria (n=52 and 51 respectively). The treated bacteria have been affected by the antibiotic and are statistically different to the untreated control (p-value ≪ 0.001). (**B**) The fluctuation distributions for untreated (n=51) and kanamycin treated (n=52) kanamycin resistant bacteria. The distributions differ slightly, but the difference is far lower than for the susceptible strains (p-value=0.015). The resistant strain shows very little effect from the antibiotic treatment after 30 minutes incubation. (**C**) The fluctuation distributions for untreated (n=50) and trimethoprim treated (n=50) susceptible bacteria, showing a slight decrease on average in the fluctuation levels for the treated sample. The difference in the distributions is, however, statistically significant (p-value < 0.001). (**D**) The individual measurements of the fluctuation levels of the different bacteria in the samples shown in c plotted at the time the measurement was taken along with the mean fluctuation level (dashed lines). The fluctuation levels of the trimethoprim treated bacteria decrease on average due to the continued action of the antibiotic, as shown by the linear fit (solid lines). The data in this figure show the effectiveness of our technique as a susceptibility test for distinguishing between susceptible and resistant bacterial samples for both bacteriostatic and bactericidal antibiotics. **Figure supplement 1.** Fluctuation distributions for untreated and kanamycin treated susceptible and resistant bacteria. The samples were prepared identically to those in Figure 5. (**A**) The distributions for the kanamycin susceptible bacteria. In agreement with the data in Figure 5A, the untreated (n=52) and treated (n=50) kanamycin susceptible samples are statistically different with a p-value < 0.001. The means of the distributions are 0.13 +/− 0.03 and 0.06 +/− 0.02 for the untreated and kanamycin treated samples respectively. (**B**) The distributions for the kanamycin resistant bacteria. In agreement with the data in Figure 5B, the untreated (n=50) and treated (n=50) kanamycin resistant samples are not statistically different with a p-value of 0.8. The means of the distributions are 0.2 +/− 0.03 and 0.2 +/− 0.04 for the untreated and kanamycin treated samples respectively. **Figure supplement 2.** Additional data for trimethoprim experiments. (**A**) Fluctuation distributions for susceptible bacteria that are untreated (n=51) and treated with trimethoprim (n=54) as for Figure 5C and D. The distributions are statistically different (p-value < 0.001) and have means of 0.15 +/− 0.04 and 0.14 +/− 0.03 for the untreated and trimethoprim treated samples respectively. (**B**) The individual measurements for a plotted at the time they were taken. A decrease is observed for the trimethoprim treated sample, as was observed in Fig. 5D. (**C**) Fluctuation distributions for an untreated sample (n=52) and a sample that was incubated in trimethoprim for 30 mins, as for the data in a, but resuspended and imaged in LB without the antibiotic (n=49). The mean of the distribution for the control is 0.15 +/− 0.04 and the mean for the trimethoprim treated sample imaged in LB is 0.14 +/− 0.03. The distributions are not significantly different (p=0.5), indicating that the effects of trimethoprim can only be observed if the antibiotic remains in the solution if the pre-incubation period is half an hour. The fluctuation distributions are statistically similar (p = 0.5). (**D**) The individual measurements for c plotted in time. No decrease in the fluctuation values for the samples are observed. Therefore, the samples need to remain in trimethoprim while imaging in order to observe the effects of this bacteriostatic antibiotic on the fluctuations.

It could be argued that the effect of a bacteriostatic antimicrobial may not be detectable by SCFI or other fluctuation-based AST methods as this class of antimicrobials does not kill the bacteria but only stops them from replicating. However, the data reported in Figure 4B demonstrate that SCFI is not a binary viability assay, and does not require the bacteria to be dead to see a difference in their metabolic activity. Accordingly, we tested the use of SCFI to detect the effects of the bacteriostatic antimicrobial trimethoprim. This is a commonly prescribed antimicrobial for Urinary Tract Infections (UTIs) to which there is widespread resistance (ESP, 2017). The antimicrobial inhibits the reduction of dihydrofolic acid to tetrahydrofolic acid, which is necessary for DNA synthesis. After a 30-minute incubation of trimethoprim susceptible E.coli bacteria with 20 *μ*g/ml trimethoprim at 37 degrees in a shaker, the bacteria were deposited on a glass coverslip using a buffer of LB also containing 20 *μ*g/ml trimethoprim. A control sample that was incubated in LB without trimethoprim was also analysed. The sub-bacterial fluctuations of the bacteria in the samples were measured (Figure 5C and Figure 5 - figure supplement 2A). The distribution for the treated sample in Figure 5C manifests a clear reduction overall compared to the untreated control sample (p-value < 0.001 with mean values of 0.22 ± 0.03 and 0.25 ± 0.05 respectively. The fluctuation level of each bacterium was plotted against the time the measurement was taken (Figure 5D and Figure 5 - figure supplement 2B) and shows that the fluctuations reduce over time due to the continued action of the antibiotic. When the assay was repeated without the inclusion of trimethoprim in the deposition and imaging buffer, there was no difference in fluctuation between treated and untreated samples and the fluctuation values did not decrease over the measurement time (Figure 5 - figure supplement 2C and D). This confirms that SCFI is specifically measuring the effect of trimethoprim as it is well established that removal of a bacteriostatic drug enables the bacteria to return to their normal metabolic activity.

Figure 5D clearly demonstrates how SCFI can provide real-time monitoring of bacterial samples as they are exposed to an antimicrobial. To illustrate this further, we recorded the fluctuations of a single bacterium as it was treated with 500 *μ*g/ml of the bactericidal antibiotic polymyxin B (video 3) in a flow chamber. This is a fast acting antimicrobial and has been observed to reduce bacterial fluctuations in seconds using surface plasmon resonance imaging (Syal et al., 2015). Using SCFI, we could directly observe where and for how long these fluctuations occur. The fluctuation level was measured for a bacterium in LB before adding polymyxin B (time = 0) while continuously measuring its fluctuations. The standard deviation of each 20s section of the recording was found (Figure 6). A control sample was also measured in the same way where LB buffer was flowed through instead of polymyxin B (video 4). The fluctuations subside to basal levels (equivalent to the dead bacteria in Figure 4A) after approx. 6 min. The high fluctuation values while adding the antibiotic (time = 0) are the effects of the flow in the sample chamber. We record a linear decrease of the fluctuations after injection of the antibiotic that contrasts with the sharp decrease observed in the plasmon resonance data (Syal et al., 2015). The reason for this discrepancy is not obvious but it may be due to the fact that the two methods measured different aspects of the cellular activity. It should be noted that SCFI can clearly observe movement at the sub-bacterium level while the cell as a whole remains still. Specifically, SCFI allows monitoring of the cellular activity independently of Brownian motion of the bacterium.

**Figure 6.**
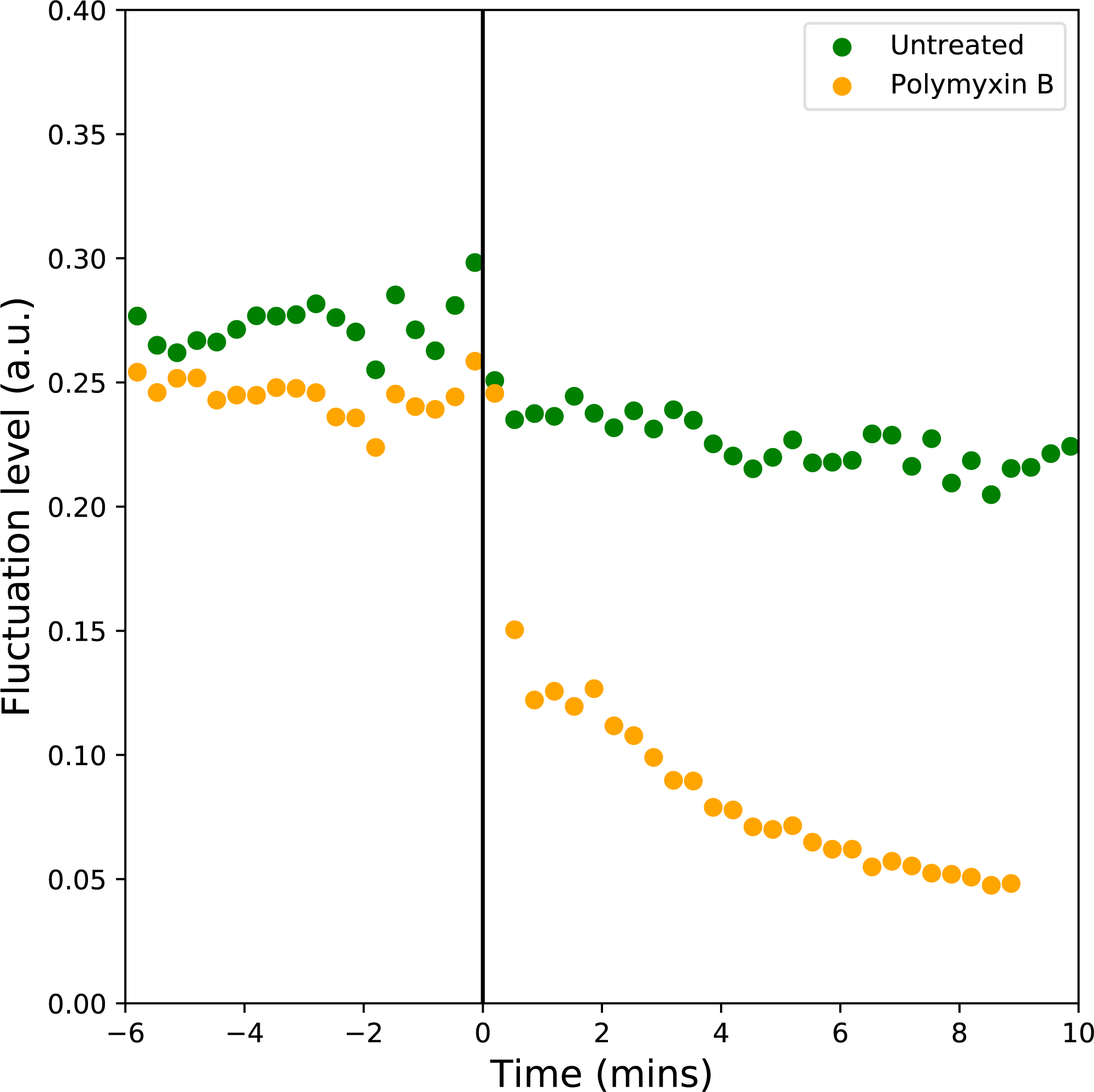
Real-time observation of the effects of polymyxin B on an *E. coli* bacterium. Plot showing the normalised standard deviation in intensity, calculated as for the data in Figs. 3 and 4, for 20s sections of videos of single bacteria in LB. At time=0, polymyxin B 500 *μ*g/ml in LB is added to one sample (polymyxin B), while LB alone was added to the other (untreated). The fluctuation level of the untreated sample remains constant, whereas the fluctuation level of the Polymyxin B treated sample rapidly decreases over 6 minutes before it plateaus as the bacterium is dying. A linear line of best fit is fitted to the data from time=0 to 6s to illustrate this, excluding outliers that have very high fluctuation levels due to the addition of the fluid. The videos of the polymyxin B treated and untreated bacteria are shown in videos 3 and 4 respectively. **Figure supplement 1.** Bright-f eld image of a Bacillus subtilis bacterium. This image corresponds to the scattered light video of the Bascillus subtilis bacterium in video 5. Scale bar is 1*μ*m.

Finally, in order to assess the range of applications of SCFI, it is necessary to consider the important class of Gram-positive bacteria. Due to their thicker cell envelope they could present smaller fluctuations and be more difficult to study by this methodology. However, we confirmed that SCFI can detect fluctuations in Gram-positive bacteria similarly to those observed in the Gram-negative E. coli. Video 5 shows a Bacillus subtilis bacterium exhibiting fluctuations and a corresponding optical image was also taken (Figure 6 - supplementary figure 1).

## Discussion

The rapidity and simplicity of SCFI combined with the depth of details contained in the fluctuation distributions are the most important benefits of this technique. AST of bacterial samples using SCFI takes tens of minutes rather than tens of hours, with further time savings potentially achievable using automation. From the examples shown here, it becomes apparent that the direct observation of sub-bacterial fluctuations using SCFI opens up the attractive possibility of examining different bacterial fitness or activity states within a population, as well as the mode of action of different antimicrobials. We have shown that each distribution provides a detailed fingerprint of the state of a bacterial population that can be immediately correlated to the effects of antimicrobials or of other induced metabolic stresses. Furthermore, the single-cell level of the measurements allows the investigation of bacteria in unusual growth phases, for example, persister cells, which are metabolically inert but viable and are increasingly being seen as important for recurrent infections recalcitrant to antibiotic therapy. More generally, SCFI could also be applicable to assessing the activity levels of other cell types, as they may manifest similar vibrational pattern (Biswas et al., 2017).

Bacterial fluctuations are an alternative phenotypic characteristic to growth and their measurement by SCFI enables more accurate and rapid characterisation of bacterial samples. The establishment of an absolute scale for bacterial fluctuations is a critical milestone in this new area of research and will have fundamental implications in the definition of a new AST. The SCFI method possesses unique characteristics that enable the rigorous definition of this measuring scale. One of the most important characteristics of any method to detect bacterial fluctuations is that it should not to be dependent on the specific instrument design. To test that SCFI satisfies this condition we compared the results to those take using a second microscope using a shorter laser wavelength, a different video camera, a smaller magnification and larger illumination field. We can confirm that the two instruments, when observing the fluctuation levels of a bacterial sample, produce results that are statistically equivalent (Figure 7).

**Figure 7.**
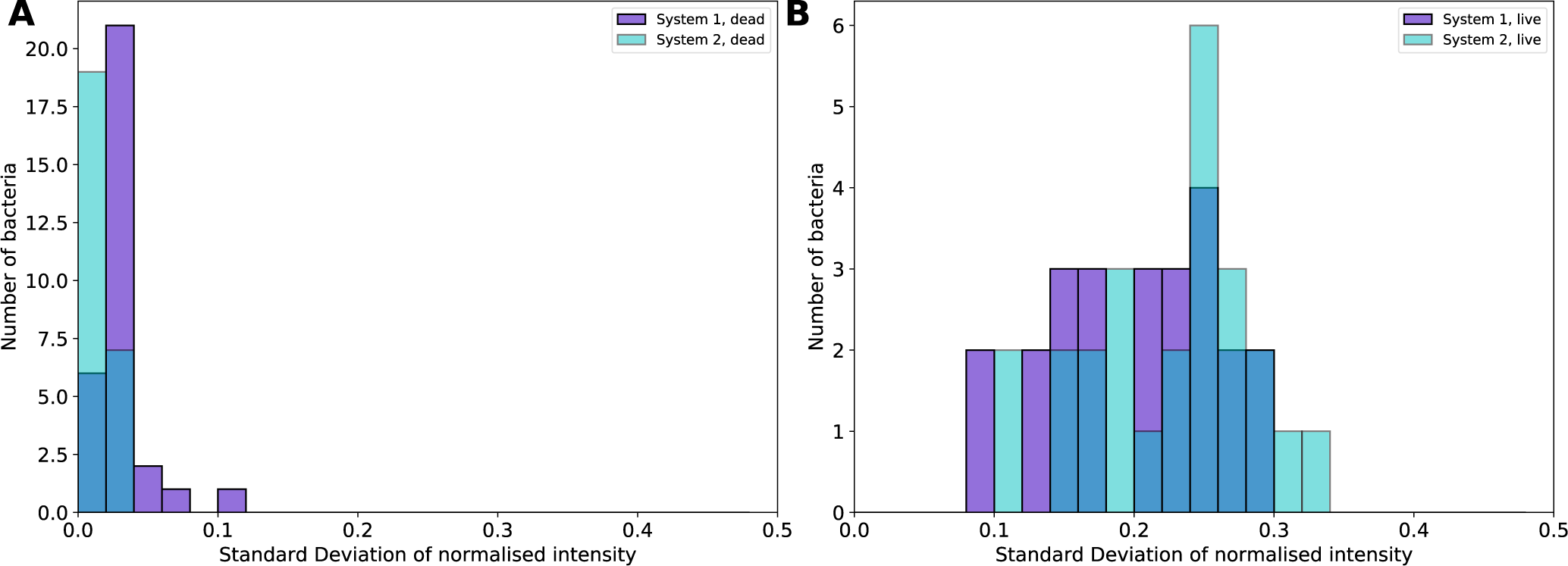
(**A**) Distribution of fluctuations levels for bacteria treated with formaldehyde (n=20) and measured on two different SCFI systems show no statistically significant difference. System 1 has been described in the previous part of the paper, while system 2 makes use of an sCMOS camera with lower sensitivity, a smaller pixel size and a 488 nm laser beam with a 5x larger beam diameter. (**B**) Similar comparison as in **A** using exponentially growing bacteria (n=25). Also in this case there is no statistically significant difference between the results produced by the two instruments.

Under the highly controllable experimental conditions described in the present work, we have shown how SCFI allows a direct comparison between untreated and treated bacterial samples for both bactericidal and bacteriostatic antibiotics. We further demonstrate that bacterial samples that are in different metabolic states can be discriminated. By revealing the subcellular level of bacterial fluctuations, SCFI offers a clear path towards the definition of a rapid, simple and rigorous protocol to define the state of the population in a bacterial sample with countless innovative applications from healthcare to biotechnology.

## Methods and Materials

### Bacterial preparation

The bacterial strains were stored at −80 using cryo-protect beads. For each week of experiments, an overnight culture was grown either from a single colony from a previous LB agar plate-culture or from the frozen stock, and was plated on LB agar. The plate was incubated at 37 °C overnight and stored at 4 °C for up to a week.

For each day of experiments, and overnight culture was prepared by suspending a single colony from the LB agar plate in 10 ml LB broth and incubating in a 37 °C shaker (120 rpm in a sealed 50 ml falcon tube). A fresh subculture was prepared for each experiment so that the bacteria can be grown to be in the same state in order to accurately compare between samples. We verified that the overnight culture remains viable throughout the day by plating 100 *μ*l of a 10^−6^ dilution of the overnight culture on agar at different points throughout the day. The viability remained unchanged with colony counts of 104, 99 and 163 for plates made up at 10am, 2pm and 6pm.

DH1 *E. coli* was used for most of the experiments. An *E. coli* K-12 strain containing a pUC-derived pK18 vector conferring resistance to kanamycin was used for kanamycin resistance experiments. The Gram-positive bacterium used was Bacillus subtilis, subsp. subtilis str. 168. The antibiotics used were kanamycin sulphate (Biobasic, KB0286) from a stock solution of 100 mg/ml in PBS, trimethoprim (Sigma T-7883) from a stock solution of 20 mg/ml in DMSO and polymyxin B (Sigma P-1004) from a stock solution of 50 mg/ml in purified water.

For the dead sample (Figure 4), the cells were chemically fixed such that they were killed but remained intact. A subculture in LB of an overnight culture was grown to an OD600 of approximately 1 then 1 ml of this solution was then rinsed by resuspending twice in 1 ml PBS (centrifuging for 3 min at 8K rpm) then resuspended and incubated in 200 *μ*l of 2% formaldehyde in PBS for 1 hour. 1 ml PBS is then added to the solution, which is rinsed twice by resuspending in 1 ml PBS and finally resuspended in 1 ml 0.06% sodium azide in PBS. The sample is imaged in PBS. A sample of dead bacteria was plated on LB agar and no growth was observed after incubation at 37 °C overnight, confirming that all the bacteria in the sample have been killed.

For the stationary state sample (Figure 4), a sample was taken directly from the overnight culture. A further sample from the overnight culture was spun down and the supernatant used to dilute the first sample to a concentration suitable for imaging. The supernatant was also used as the buffer to flow through when binding the bacteria to the sample surface. This ensures that the bacteria remain in stationary state by ensuring the dilution and sample preparation buffer has the same chemical composition as the sample.

The exponential state bacteria (Figure 4) were prepared by incubating a subculture from the overnight sample (10 ml total of a 1/100 dilution of the overnight culture in LB) until it reached an OD600 that was known to correspond to the bacteria being in an exponential state (Figure 4 - figure supplement 1). Fresh LB was used as a buffer for imaging.

For each sample in the antibiotic experiments (Figure 5), a subculture containing 30-50 *μ*l of the overnight culture (depending on the growth speed of the strain) in 10 ml fresh LB broth was prepared by incubating in the 37 °C shaker for approx. 1 hour 45 mins until it has reached exponential phase, as confirmed by OD600 measurements. Antibiotics were then added to the relevant samples and both the treated and untreated samples were incubated for a further 30 minutes in the 37 °C shaker. The solutions were then centrifuged (8K rpm, 3 minutes) at room temperature and the bacterial pellet resuspended for the experiments. The kanamycin treated samples were resuspended in LB in order to observe the effect that half an hour incubation in the solution had on the bacteria. The trimethoprim treated samples were resuspended in LB containing the same concentration of trimethoprim as the incubated sample, as the effect of the antibiotic is only observable when the antibiotic is present. The final density of bacteria in the samples is approx. 5 × 10^8^ colony forming units (cfu)/ml. The untreated control samples are always recorded using SCFI on the same day as the treated samples as the state of the bacteria can vary between days, possibly due to different characteristics of the originating colony or different buffer conditions. As microbiological controls, the OD600 of each sample was measured before and after incubation with/without antibiotic to assess growth and check that the OD600 does not exceed the level for exponential growth. In addition, each sample was plated on LB agar after incubation to assess the viability by counting colonies on the plate after overnight growth at 37 °C. If the growth was not confluent (con), the number of colonies was counted and compared to the cfu/ml of the sample, calculated from the OD600, in order to quantify the number of bacteria that remain viable. This provides a check of the state of the samples that were analysed. The OD600 before and after incubation and the plating results for all samples in the main text are shown in Supplementary File 1.

### Preparation of the flow cells

The samples were analysed in functionalised glass bottomed petri dishes containing a flow cell. The purpose of the functionalisation is to coat the glass with amino functional groups (NH_2_), to which the antibodies can covalently bind, creating an even and dense coating of antibodies on the surface, to which the bacteria can bind. The procedure use is based on a previously described ethanolamine procedure (Ebner et al., 2007).

6g of ethanolamine hydrochloride (Sigma Aldrich, E6133) is dissolved in 16 ml DMSO (Honeywell Research Chemicals, 41640) by heating the solution in a vial at 65 °C in an oil bath on a hot plate for 20-30 mins, using a magnetic stirrer to agitate the solution. Once the solution has cooled to room temperature, a layer of molecular sieves (0.4 nm, 1/16 inch pellets, Merck, 105743) is added to the vial and it is placed, with the lid removed, in a vacuum chamber to degass the solution for 30 mins. Approx. 2 ml of the solution is pipetted into each glass bottomed petri dish (Cellview cellculture dish, Grenier Bio-One, 627861), the dishes are covered and incubated overnight. The petri dishes are then rinsed with DMSO followed by ethanol and dried in a stream of nitrogen. The dishes can be stored for a few weeks in a desiccator until use.

The flow cell is constructed by adhering a small coverslip to the glass surface of the petri dish using two parallel strips of double sided tape. The coverslips are cleaned in ethanol, followed by distilled water and dried in a stream of nitrogen prior to constructing the flow cell. Antibodies, samples and buffers are pipetted into the flow cell from one end and liquid is drawn through using filter paper at the opposing end.

### SCFI instrumentation

Our technology uses label-free high-resolution detection of scattered light from the bacteria to image the nano-fluctuations in real-time on a high resolution camera. In order to detect the fluctuations, the bacteria are aligned and bound to a surface above which an evanescent wave is generated using a laser (1mw, 561 nm) such that only a ~100 nm deep portion of the lower section of the bacteria is illuminated.

The evanescent field is generated above the glass coverslip by total internal reflection of a laser beam using a high numerical aperture (NA) objective lens. A piezo inertia actuator provides movement of the sample in the vertical direction (z-direction) in order adjust the focus position. The reflected beam from the surface is deflected away from the optical detection system. The scattered light from objects located within the evanescent field is collected through the high NA lens and reflected to pass through a tube lens positioned such that the image plane is located on the sensor of a high resolution CMOS camera. The sample is mounted on an x-y slip-stick positioning stage in order to position objects in the evanescent field for imaging with 1*μ*m accuracy. The camera is capable of detecting lateral movement of 30 nm. Vertical movements of a few nanometres of scattering objects within the evanescent field result in changes in the intensity of the scattered light on the camera, due to the rapid exponential decay in intensity of the evanescent field away from the sample surface.

Two systems with slightly different components were compared in Figure 7. System 1, as described above, was used for the majority of the results presented in the paper. It uses the following components:

1. Hamamatsu Orca4 sCMOS (pixel size 6.5 *μ*m × 6.5*μ*m, quantum efficiency 70% at 561 nm, readout noise 1.5 *e*^−1^ (RMS)).
2. 561 nm fibre-coupled diode laser (Vortan, Stradus). The beam diameter after collimation is 1 mm. Typical power after the fibre 1 mW.
3. Nikon TIRF 1.49 NA objective lens
4. Image size on the video camera sensor 26.5 nm/pixel

System 2 has a similar design to System 1 but the tube lens is mounted directly after the Nikon objective lens, the magnification is smaller and after collimation the laser beam expansion is 5× larger than in System 1. System 2 uses the following components:

1. Thorlabs Quantalux sCMOS (pixel size 5.04 *μ*m × 5.04 *μ*m, quantum efficiency 55% at 488 nm, readout noise <1.5 *e*^−1^ (RMS)).
2. 488 nm fibre-coupled diode laser (Vortan, Stradus). The beam diameter after collimation is 5 mm. Typical power after the fibre 30 mW.
3. Nikon TIRF 1.49 NA objective lens.
4. Image size on the video camera sensor 43 nm/pixel.

When measuring the same bacterial sample in the same temperature conditions using the two SCFI systems there is no statistically significant difference between the two distributions. This test was repeated for dead and live bacterial samples and shows a remarkable agreement between the two systems. We can conclude that SCFI is an extremely robust way of measuring the scattered signal from a bacterium and the mean fluctuation value returned is not strongly dependent on the commercial components used. We have preliminary data showing that the angle of collection is more critical. This means that using an objective lens with a smaller NA (i.e. Nikon TIRF 1.42 object lens) reduces the actual value of the mean fluctuations. It is not unreasonable to think that normalising the data by the solid angle of collection used in a particular SCFI system may reduce the discrepancy between different systems in a similar way as the normalisation by pixel intensity used in our analysis software reduces the variability between different bacteria.

### Experimental protocol

To firmly bind bacteria to the functionalised glass surface in the flow cell, immediately prior to imaging a solution containing appropriate antibodies is incubated in the flow cell for 5 minutes. For the *E. coli* experiments, anti-E. *coli* antibodies (abcam ab137967) were used and for Bacillus subtilis, anti-Gram positive antibodies were used (abcam ab20344).

The bacterial sample is then flowed into the flow cell and incubated for approximately 10 minutes with occasional gentle flow to allow time for antibody capture of the bacteria. LB is then flowed through, removing unbound bacteria. The bound bacteria are largely aligned with the direction of the flow. This results in more consistent data than randomly oriented bacteria, as the bacteria have different scattering characteristics depending on their alignment with the evanescent f eld due to their elongated shape. The bacteria can be observed optically on the high resolution camera using bright-field illumination with a field of view of approx. 60×60 *μ*m^2^.

Bacteria are individually positioned in the centre of the evanescent field. Videos (20 s long, 20 frames/s) are taken of each bacterium to record the intensity fluctuations. An optical image of each bacterium is also recorded. We ensure that each bacterium is fully bound to the surface such that no movement of the whole cell can be seen in either the bright-field or scattered light view and that it is well aligned in the evanescent f eld. The fluctuation levels of approximately 50 bacteria are recorded for each sample to determine the produce the fluctuation distributions.

### Fluctuation analysis method

The bright-field image of a bacterium is used to locate the coordinates of the central point of the bacterium. These coordinates are then used to position a 30×30 pixel analysis area on the centre of the bacterium for each frame in the scattered light video.

Pixel intensities in the analysis area are extracted from each frame and normalised by the average intensity of the analysis region through the duration of the video. This normalisation compensates for changes in exposure, laser intensity and scattering intensity. Pixels saturated throughout the video are removed from the analysis. The standard deviation of the intensity of each pixel in the analysis area is calculated through time. A bacterium’s fluctuation level is defined as the average of the pixel standard deviations over the analysis region. This process is illustrated in Figure 4 - figure supplement 2.

### Simulations

Simulations in Figure 1B were performed using a 2-dimensional finite-difference time-domain (FDTD) method. For this, the MIT Electromagnetic Equation Propagation (MEEP) FDTD system was used (Oskooi et al. (2010)). The beam was sourced with a Gaussian-prof le, and the refractive index of the glass, water and the bacterium are 1.515,1.33 and 1.38 respectively. The incident angle of the laser is 62°, the wavelength used in this case is 561 nm.

Further modelling of the scattering interaction between a bacterium and the light field (Figure 2 - figure supplement 1), was carried out using a finite element solver (COMSOL). The bacterium is modelled as a uniform material with refractive index of 1.4 lying on a glass substrate (*n* = 1.5) and suspended in water (*n* = 1.33). A Gaussian beam (polarised out-of-plane *E*_*z*_) with beam-width 8 *μ*m is incident from the glass side at the TIR angle 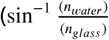 and the scattered field is calculated.

To quantify the scattering pattern, we perform two simulations, one with the bacteria having a background index (*n* = 1.33) and the second with *n* = 1.4 and subtract the two fields (this amounts to an effective background subtraction, which in the experiment occurs by the viewing angle of the lens preventing the TIR beam from being viewed). To perform this background subtraction, we output the fields from the triangular mesh onto a Cartesian grid. The plot in Figure 2 - figure supplement 1 shows the net Poynting vector 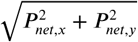 plotted as a function of position, which gives an estimate of the net power flow (defined as *P*_*net, x*_ = *P*_*bacterium, x*_ − *P*_*bkgd, x*_) and we can easily see the two peaked response (shown more clearly in the zoomed-in image and linecut), corresponding to enhanced scattering from the ends of the bacterium. We would like to note that in subtracting the Poynting vectors directly and not the field components themselves, we are assuming that the cross terms (*E*_*bacterium, z*_ × *H*_*bkgd, y*_) are small compared to (*E*_*bacterium, z*_ × *H*_*bacterium, y*_ We believe this assumption is valid because of the low field overlaps and the weak index contrasts in this platform.

### Statistical analysis of the fluctuation distributions

In order to determine whether fluctuations distributions from different samples are statistically different from each other, Welches two sided t-test for two independent samples with non equal variance was used to find the t-statistic and corresponding p-value. The degree of freedom used for calculating the p-value was (n1-1) + (n2-1) where n1 and n2 are the number of bacteria in each of the samples to be compared respectively. The level of statistical significance to reject the null hypothesis that there is no difference between the means is taken to be p=0.05. The p-values for all plots are presented in Supplementary File 2.

## Acknowledgments

This work was supported by the University of Bristol through a grant from the EPSRC Institutional Sponsorship. It was also supported by the Elizabeth Blackwell Institute through a grant from the MRC Confidence in Concept Scheme (MC_PC_16039). We also received support through a grant from the Longitude Prize Foundation. This work was funded by grants MR/N013646/1 NE/N01961X/1 and EP/M027546/1 to M.B.A. from the Antimicrobial Resistance Cross Council Initiative supported by the seven research councils. N.M.R.’s time is supported by the National Institute for Health Research (NIHR) Collaboration for Leadership in Applied Health Research and Care West (CLAHRC West) at University Hospitals Bristol NHS Foundation Trust.

C.R.B. thanks Dr Jacqueline Findlay for preparation of bacterial strains and C.R.B and M.A. thank Dr Henkjan Gersen for reviewing the manuscript. M.A., C.R.B. and I.M. thank Ariel Blocker for assistance with early stage planning of the experiments. M.A. thanks Nick and Susan Woollacott who kindly funded the equipment used in this research as well as Adrian Crimp for his essential technical support.

---

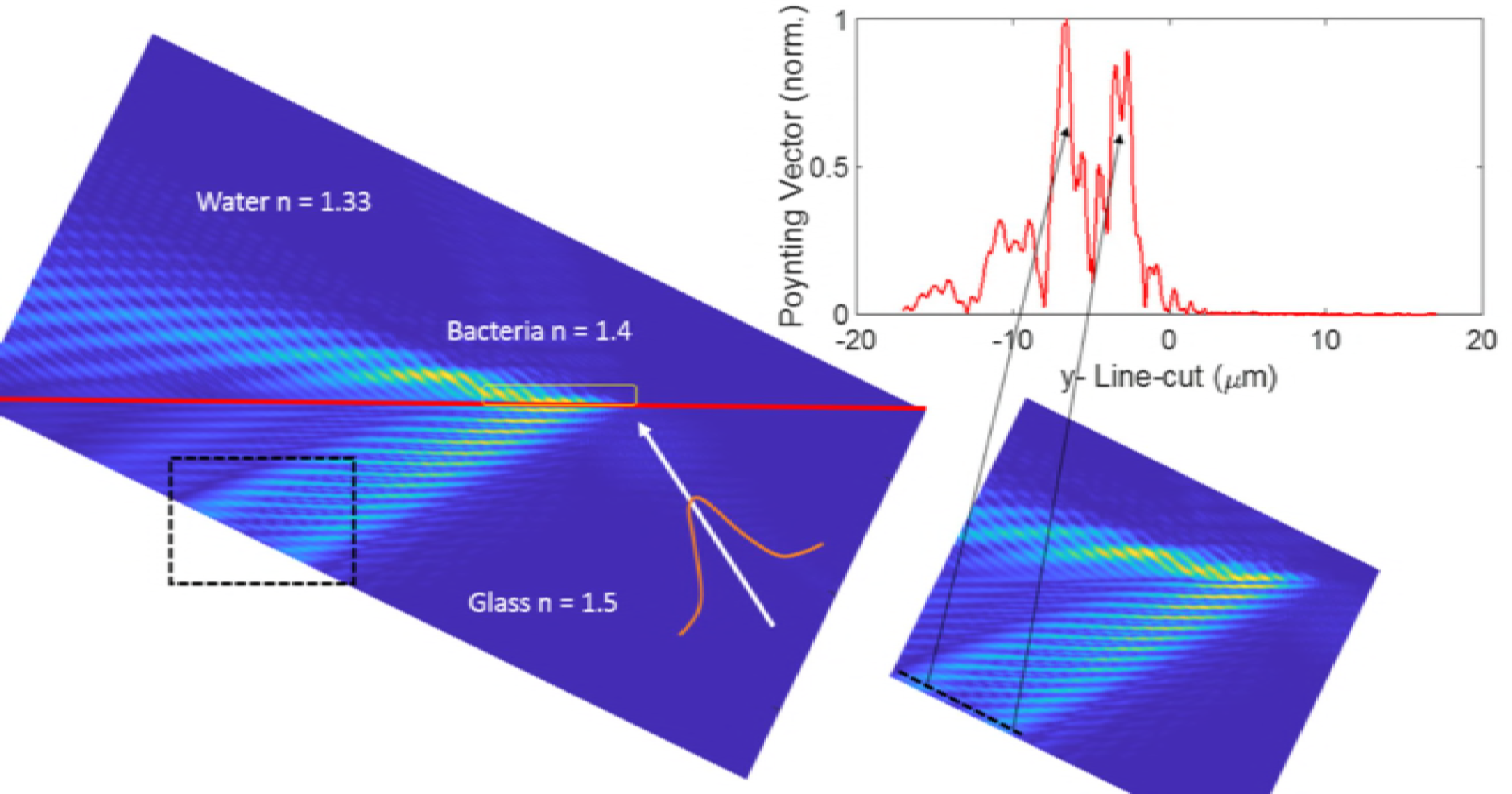

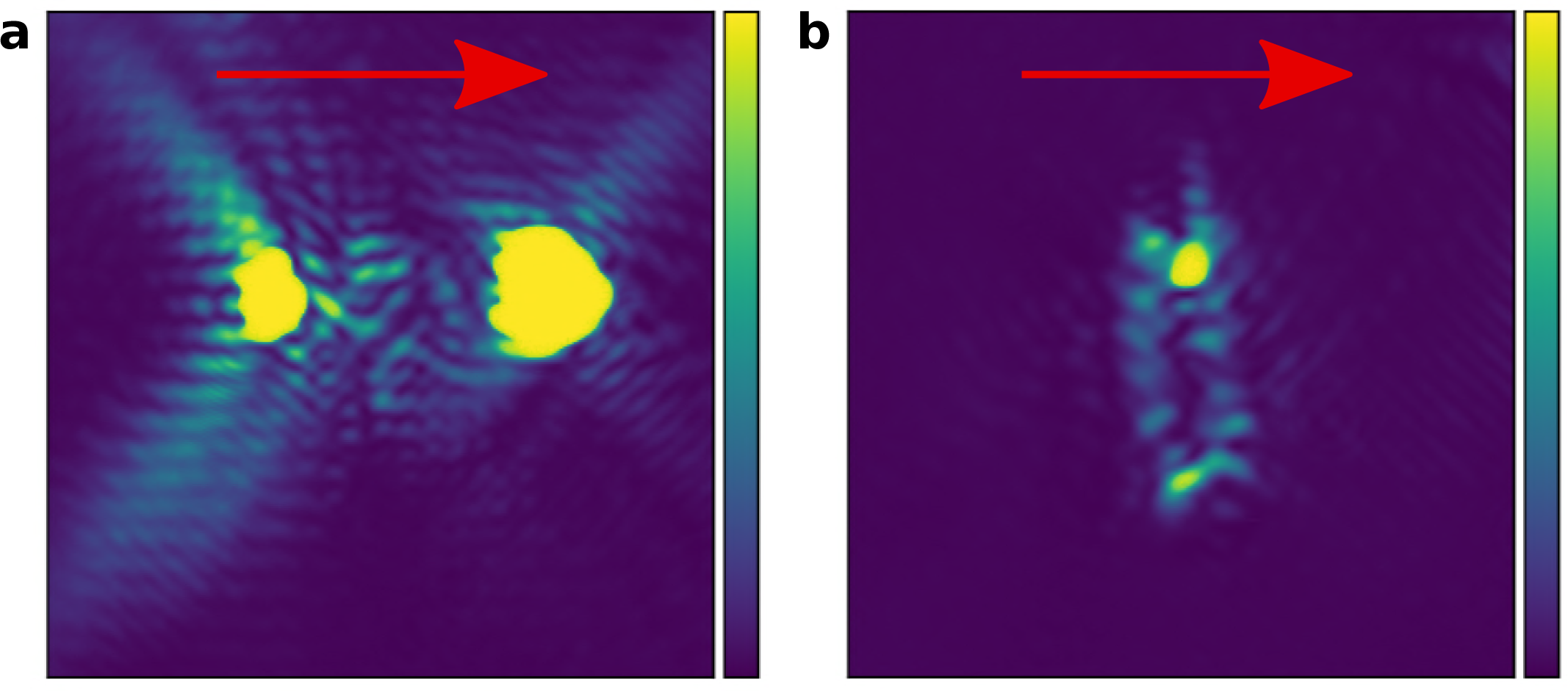

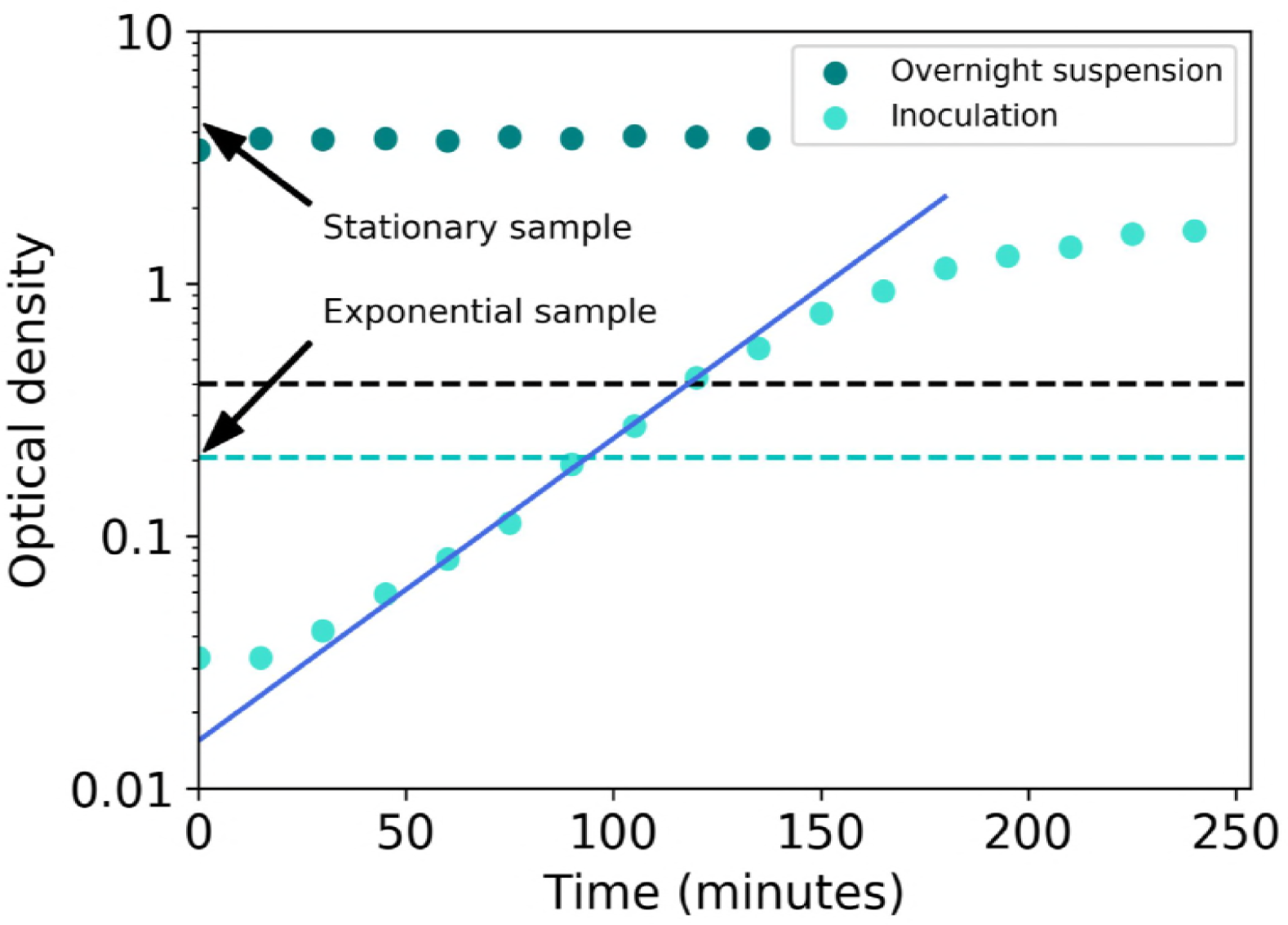

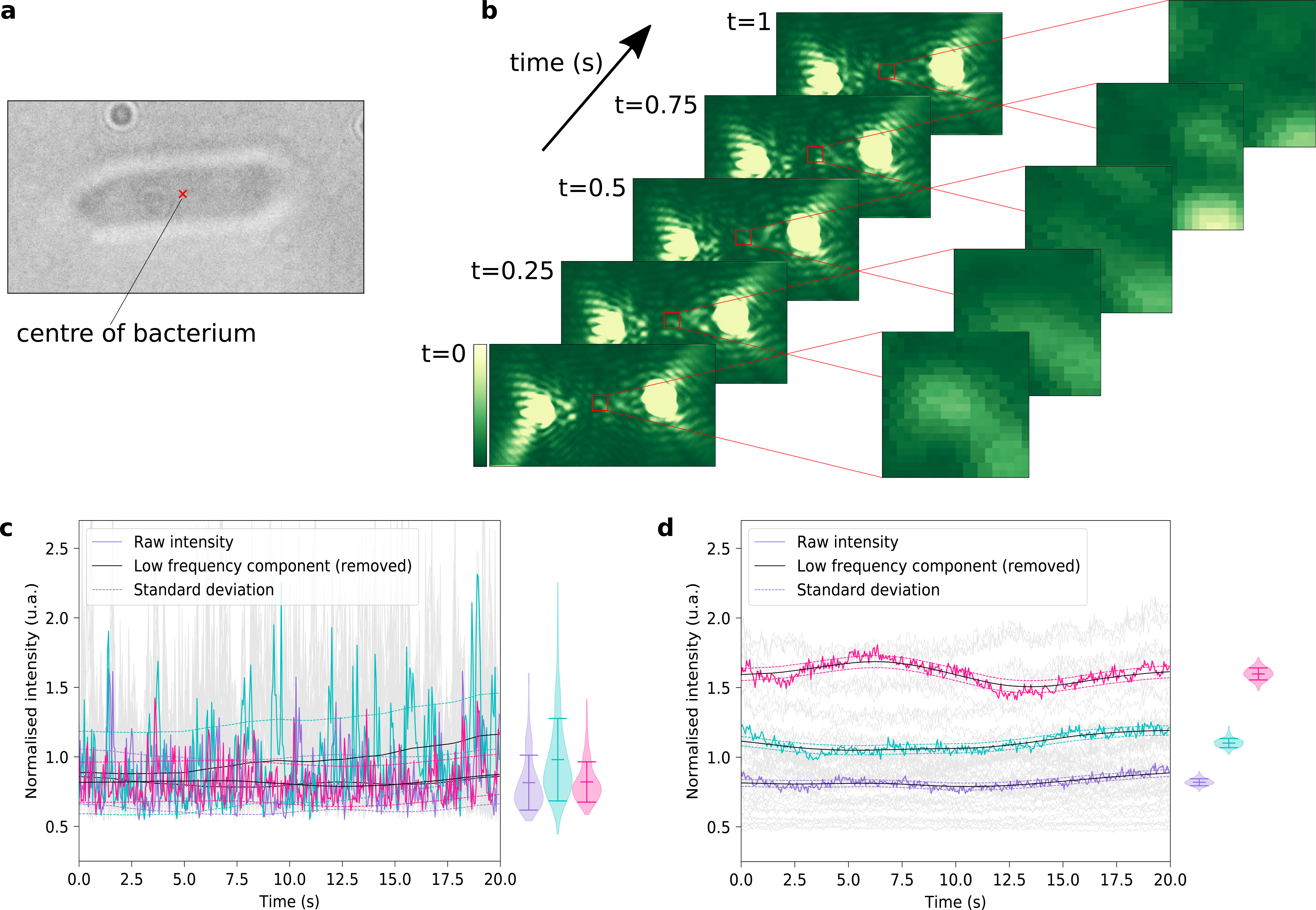

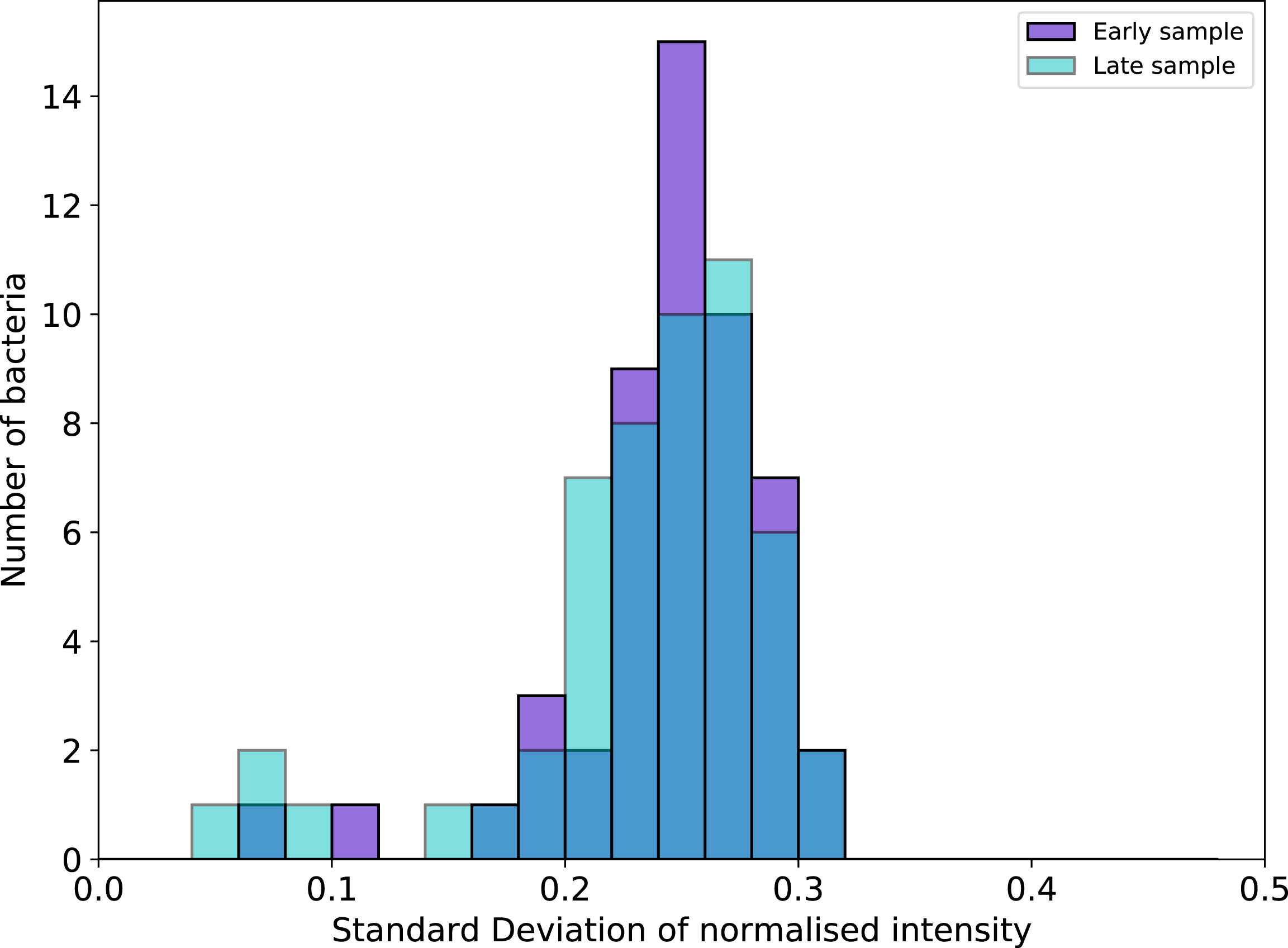

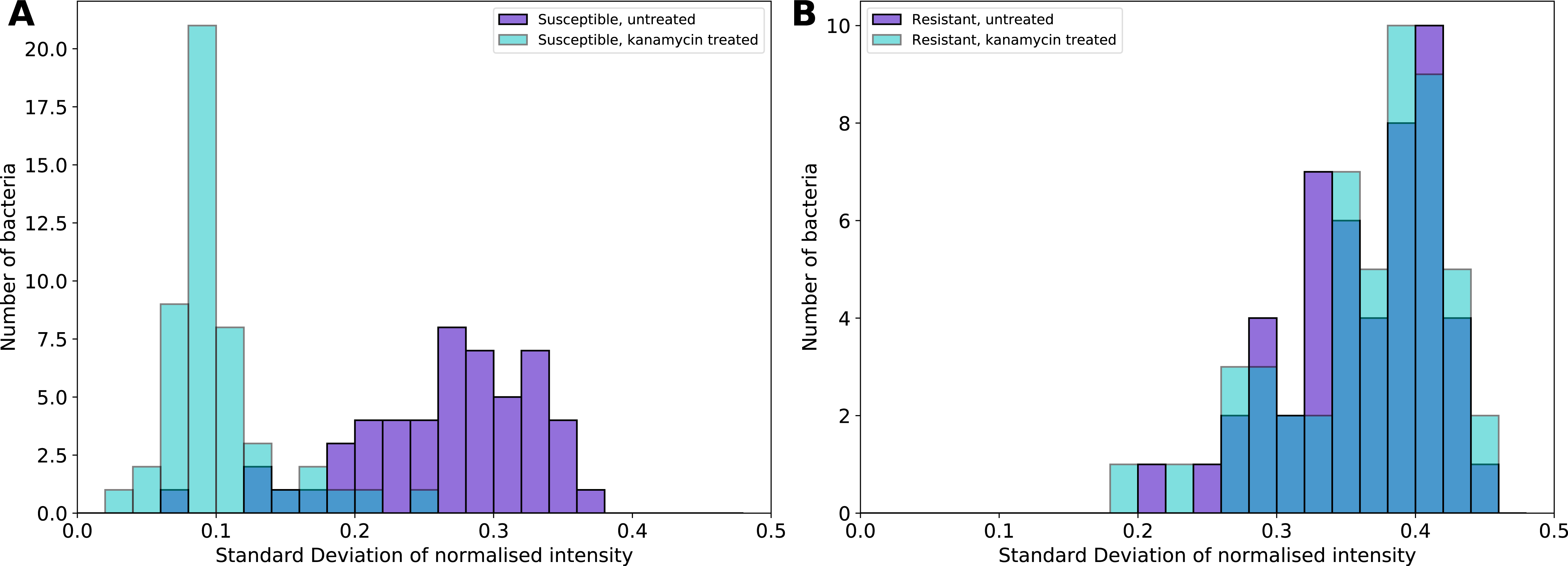

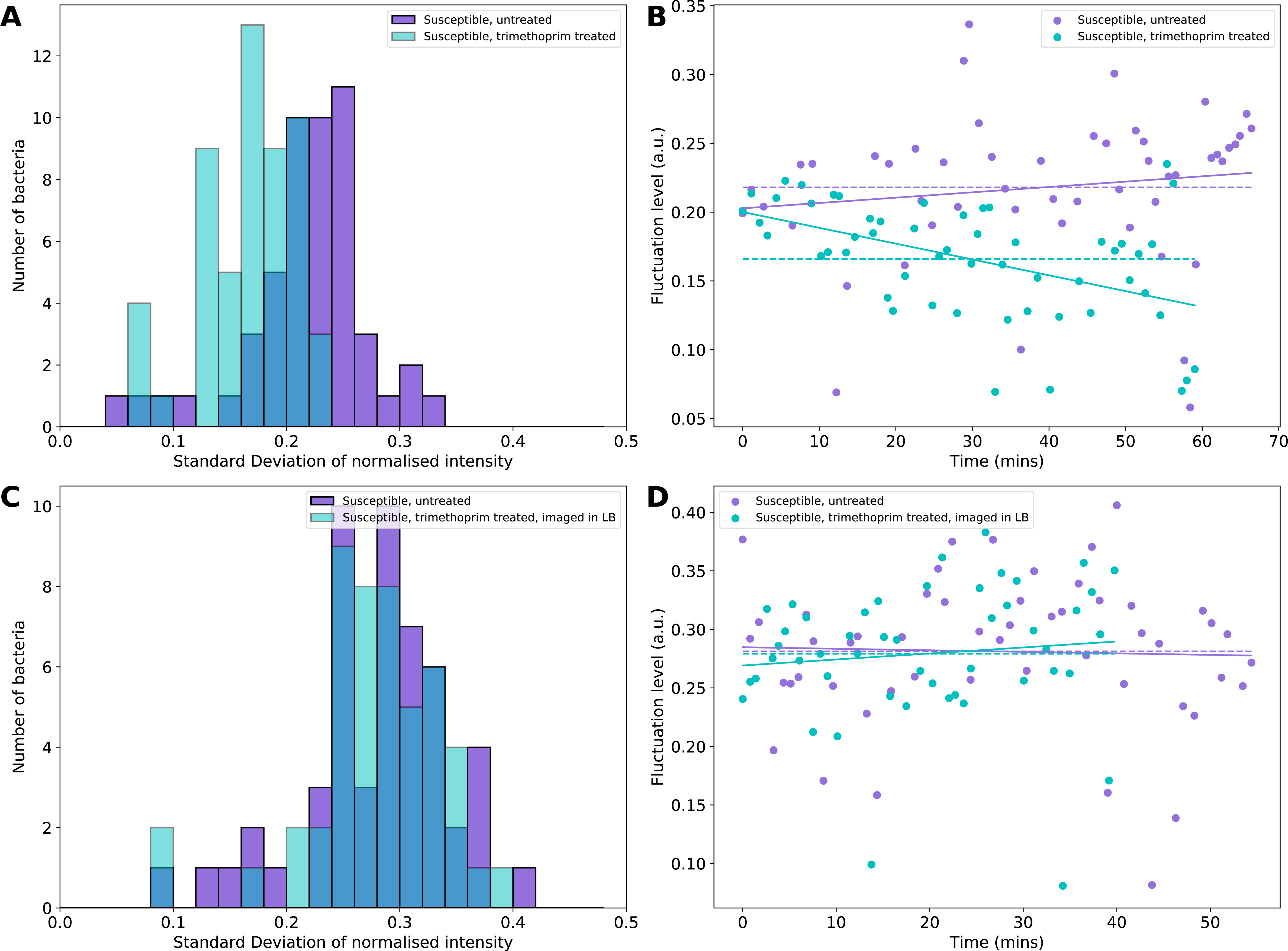

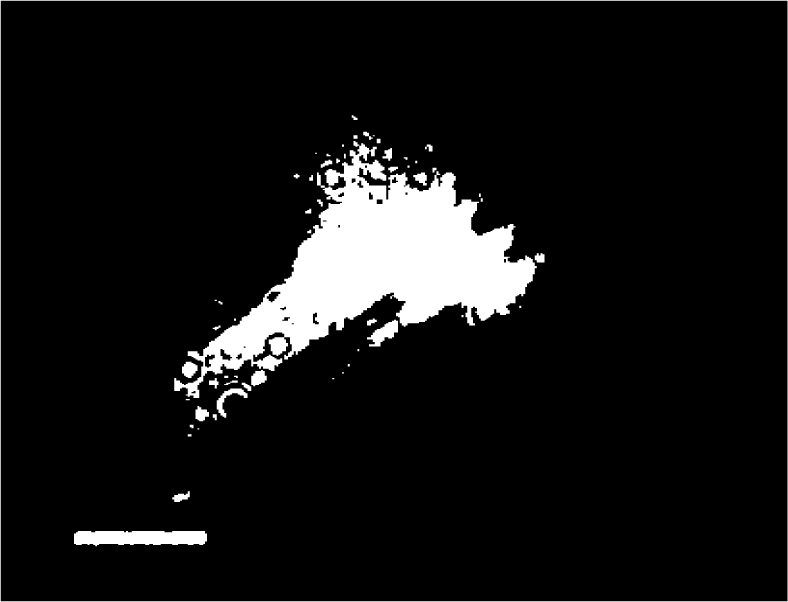

